# Nanobiopsy investigation of the subcellular mtDNA heteroplasmy in human tissues

**DOI:** 10.1101/2023.06.22.546054

**Authors:** Alexander Bury, Amy E. Vincent, Angela Pyle, Paolo Actis, Gavin Hudson

## Abstract

Mitochondrial function is critical to continued cellular vitality and is an important contributor to a growing number of human diseases. Mitochondrial dysfunction is typically heterogeneous, mediated through the clonal expansion of mitochondrial DNA (mtDNA) variants in a subset of cells in a given tissue. To date, our understanding of the dynamics of clonal expansion of mtDNA variants has been technically limited to the single cell-level. Here, we report the use of nanobiopsy for subcellular sampling from human tissue, combined with next-generation sequencing to assess subcellular mtDNA mutation load in human tissue from mitochondrial disease patients. The ability to map mitochondrial mutation loads within individual cells of diseased tissue samples will further our understanding of mitochondrial genetic diseases.

## Introduction

Inherited and somatic mitochondrial DNA (mtDNA) variation is a significant contributor to human disease (1). Inherited mtDNA variation is associated with a wide range of diseases that can present from birth to old age (2,3,4). In parallel, somatic mtDNA mutations have been shown to accumulate with age and preferentially accumulate in specific organs or tissues, and typically lead to late-onset disease such as Parkinson’s disease (PD) (5,6,7), Alzheimer’s disease (AD) (8), and pathological ageing (5,9,10). Previous work has suggested that low-level heteroplasmic variation is a common occurrence at the single cell level (11) and that the clonal expansion of these low-level variants affects mitochondrial function (11, 12) and contributes to disease (13,14,15). However, the mechanism of clonal expansion appears to vary between mtDNA point mutations and mtDNA deletions and depending on the tissue in question (16, 17, 18). Whilst some of the mechanisms, such as the clonal expansion of mtDNA point mutations in mitotic tissues, are understood (10,19). Conversely, the mechanisms driving clonal expansion of mtDNA variants in post-mitotic tissues remain elusive (19,20). The tissue and mutation specific nature of both mtDNA heteroplasmy and clonal expansion means that, to better understand these disease mechanisms, it is necessary to study mtDNA mutation within a well characterised tissue (21, 22).

Organelle, specifically mitochondrial, heterogeneity at the single-cell level is associated with localised physiological function (23,24) and can be characteristic of aberrant cellular pathways in disease (4,25).

In addition to mtDNA sequence characteristics (26), work in early developmental tissues suggests that factors such as mitochondrial locality, particularly proximity to the nucleus, influence heteroplasmy dynamics (27,28). Further studies investigating heteroplasmic variation have shown that variation can be regional, and it has been hypothesised that subcellular environment and cellular structure can impact clonal expansion (29,30). Whilst stochastic models have been used to describe the mode of clonal expansion (30,31,32, 33), they require estimates of mutation rates specific to the tissue and mutation in question and are based on limited spatial information (18,34).

Previous studies have been able to assess the expansion of mtDNA mutations longitudinally in skeletal muscle fibres and sarcomeres or, to a limited extent, able to investigate traverse patterns of mtDNA mutations across muscle fibres (29,30). However, longitudinal analysis alone cannot take into account the clonal expansion of mtDNA mutations within a branched, three-dimensional mitochondrial network and mitochondrial subpopulations can exist in foci smaller than the permitted sampling range of existing technologies (29,35). Skeletal muscle tissue is a well characterised sample medium in the investigation of clonal expansion of mtDNA mutations (29,32,36), which makes it a strong candidate sample medium for investigating clonal expansion.

The advent of nanoprobe technologies (37,38) presents a unique opportunity to study mtDNA heteroplasmy with a sufficient sampling resolution, simultaneously preserving the spatial information needed to study somatic mtDNA variation and clonal expansion. A range of micro- and nanoprobe-based technologies have been recently developed to enable the localized probing of cells and tissue (39,40,41,42,43). Our group has recently developed a subcellular nanobiopsy method based on scanning ion-conductance microscopy (SICM) to enable the subcellular sampling of mitochondria from human tissues with a spatial resolution superior to the gold standard technique, laser capture microdissection (LCM) (44).

Nanobiopsy is a form of scanning probe microscopy comprising a glass micro-or nanopipette controlled by a nanomanipulator, which can manoeuvre the pipette to subcellular regions, facilitated by fluorescence microscopy to achieve highly precise sampling of subcellular molecules (39,45). Nanobiopsy has been successfully used to study mtDNA obtained from mitochondria isolated from cultured fibroblasts (39) and human skeletal muscle tissue (44) and some of the recent applications of this technology have been recently reviewed (46).

In this study, we demonstrate that nanobiopsy is a viable tool for the study of heteroplasmic mtDNA variants in human tissue. This proof-of concept work shows that nanobiopsy can be used in conjunction with next-generation sequencing to investigate sub-cellular mtDNA heterogeneity (**Figure 1**).

**Figure 1.**
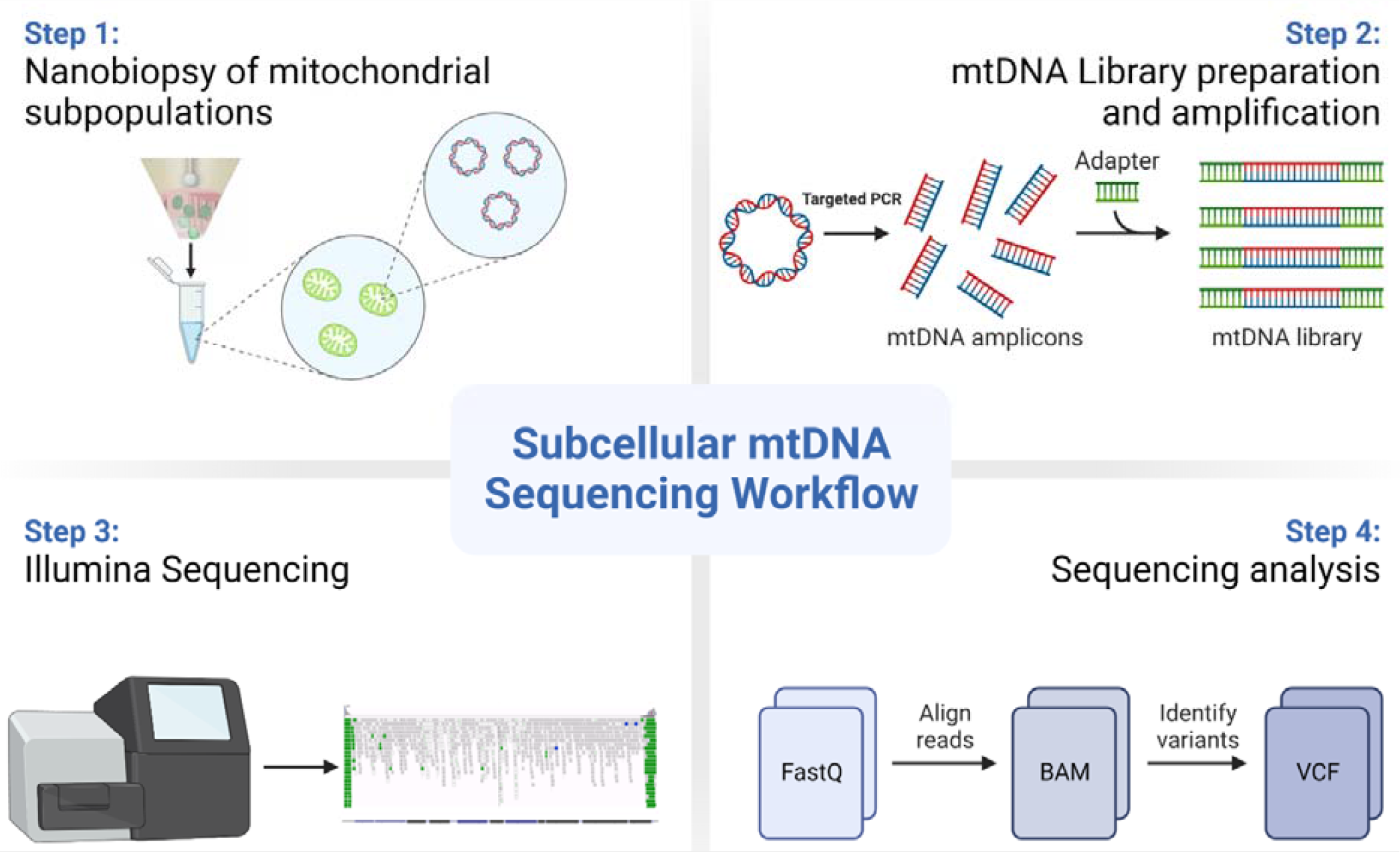
Depicted is a graphic showing the workflow for the sequencing of mtDNA isolated from subcellular mitochondrial populations obtained using nanobiopsy. Following the nanobiopsy of fluorescently labelled mitochondria from distinct subcellular foci, mitochondria are lysed and mtDNA molecules are fragmented prior to library preparation and clean-up. MiSeq sequencing was performed on the D-loop region of mtDNA and post sequencing analysis allowed alignment of reads and identification of mtDNA variants. In Step 3 the integrated genomic viewer image is adapted from Wei and colleagues (47).

## Materials and Methods

### Skeletal Muscle Tissue Biopsies

Excess skeletal muscle tissue from hamstring was collected during anterior cruciate ligament surgery with prior informed consent and ethical approval from the Newcastle Research Ethics Committee (REC: 12/NE/0267). Skeletal muscle from mitochondrial disease patients excess to diagnostic requirements was consented for research and transferred to the Newcastle Mitochondrial Research Biobank after diagnosis. Tissue was requested from the biobank and approved (REC: 16/NE/0267 - Application Ref: MRBOC ID039) and all work is performed with ethical approval from the Newcastle and North Tyneside Local Research Ethics Committee (REC: 16/NE/0267). All research was carried out with proper HTA licensing and tissue transfer agreements in place.

### Cryosectioning

Skeletal muscle tissue was cryosectioned into multiple 15μm sections. Sections were mounted on square glass microscope slides (Agar Scientific Ltd). Sections were then air-dried for 1 hour and stored at⍰− ⍰80 °C prior to staining.

### Immunofluorescent staining

Immunofluorescent staining was performed on skeletal muscle tissue sections fixed with 4% PFA, using primary antibodies raised against mitochondrial proteins: VDAC1, NDUFB8 and MT-CO1 (**Table 1**). Dehydration and rehydration of tissue sections, through a methanol gradient, was performed to permeabilise sections. Blocking and immunofluorescent staining steps were performed as previously described, with the addition of DAPI staining to allow visualisation of myonuclei (48).

**Table 1.**
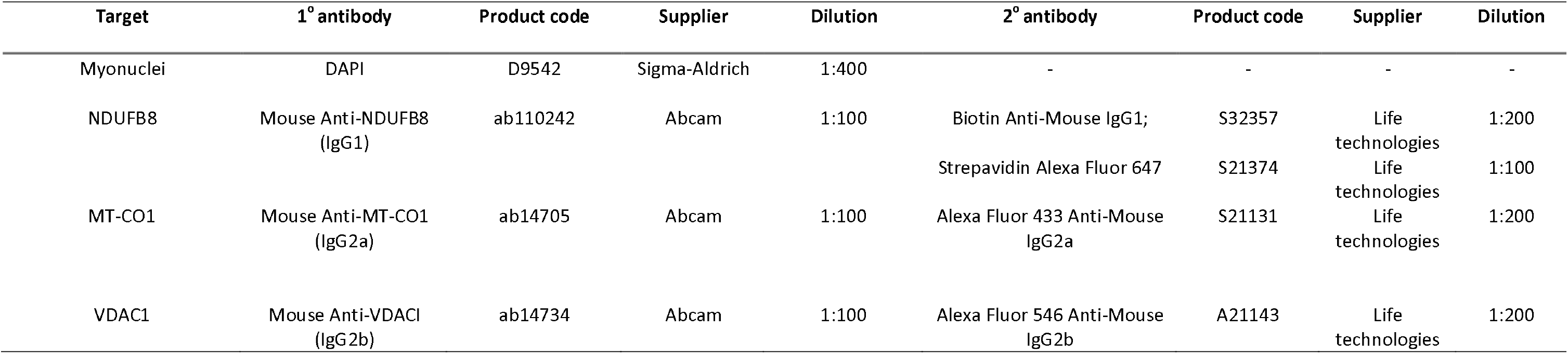
Supplementary Table 1. Quad-immunofluorescence primary and secondary antibodies. All primary and secondary antibodies are listed, used for QIF staining of specific targets in skeletal muscle tissue. The product number, supplier, dilution and antibody target are also included.

| Target | 1° antibody | Product code | Supplier | Dilution | 2° antibody | Product code | Supplier | Dilution |
| --- | --- | --- | --- | --- | --- | --- | --- | --- |
| Myonuclei | DAPI | D9542 | Sigma-Aldrich | 1:400 | - | - | - | - |
| NDUFB8 | Mouse Anti-NDUFB8 (IgG1) | ab110242 | Abcam | 1:100 | Biotin Anti-Mouse IgG1; | S32357 | Life technologies | 1:200 |
|  |  |  |  |  | Streptavidin Alexa Fluor 647 | S21374 | Life technologies | 1:100 |
| MT-CO1 | Mouse Anti-MT-CO1 (IgG2a) | ab14705 | Abcam | 1:100 | Alexa Fluor 433 Anti-Mouse IgG2a | S21131 | Life technologies | 1:200 |
| VDAC1 | Mouse Anti-VDAC1 (IgG2b) | ab14734 | Abcam | 1:100 | Alexa Fluor 546 Anti-Mouse IgG2b | A21143 | Life technologies | 1:200 |

### Sub-cellular Nanobiopsy

Nanobiopsy was performed as previously described (39,44). Micropipettes were fabricated from borosilicate glass capillaries (BF100-50-7.5, Sutter Instruments), using an SU-P2000 laser puller (World Precision Instruments). Nanobiopsy was performed using an SICM comprising an Axon MultiClamp 700B amplifier (Molecular Devices), MM-6 micropositioner (Sutter Instruments), p-753 linear actuator (Physik Instrumente), pE-300 LED illumination system (CoolLED) and an Eclipse Ti2 inverted microscope (Nikon). Control of the micropipette and electrochemical measurements were executed using the SICM (ICAPPIC Ltd.) and Axon pClamp 11 (Molecular Devices) software. The automated approach of skeletal muscle fibres was achieved with a fall distance of 2μm, fall rate of 100μm/s and a 1% current set-point.

### mtDNA isolation, enrichment, and next-generation sequencing

Briefly, the human mtDNA control region (m.1-573 and m.16024-16569) was enriched using four overlapping PCR amplicons using high fidelity TaKaRa PrimeSTAR GXL DNA polymerase (TaKaRa; **Table 2**). PCR products were visually inspected by agarose gel, pooled in equimolar concentrations, and purified using Agencourt AMPure XP beads (Beckman-Coulter, USA). Next-generation sequencing (NGS) was conducted as previously described (6,49). Briefly, pooled amplicons underwent tagmentation, amplification, Ampure-bead purification and were normalized using an Illumina Nextera XT DNA sample preparation kit (Illumina, CA, USA). Multiplex pools were sequenced using MiSeq Reagent Kit v3.0 (Illumina, CA, USA) in paired-end, 250 bp reads. Postrun data as FASTQs, limited to reads with QV ≥ 30, were exported for analysis. NGS of nanobiopsied mitochondria was compared to lysate obtained from single-cells collected by LCM (29,50) and control genomic DNA (gDNA) (Promega), sequenced using conventional sequencing methods (13).

**Table 2.**
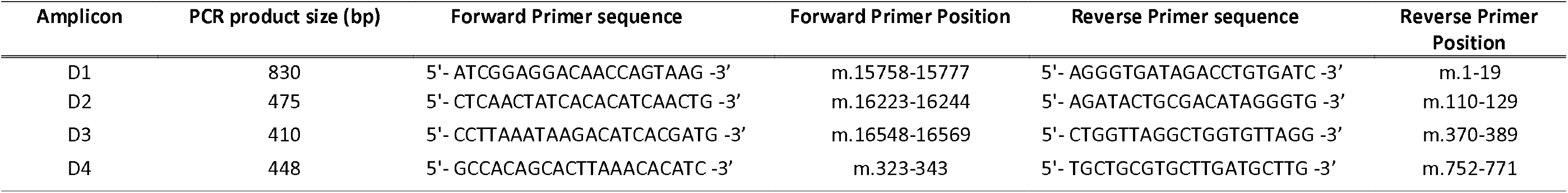
Supplementary Table 2. Targeted PCR primers. PCR product size and primer sequences (and positions) for both forward and reverse primers. D1-D4 correspond to amplicons spanning the entirety of the mtDNA control region. Primer positions are in reference to the revised Cambridge reference sequence (NC_012920.1)

### Bioinformatics analysis

Informatics analysis of FASTQ files was performed as previously described (51). Raw reads were assessed and filtered using FASTQC (v0.11.7; Babraham Bioinformatics, Cambridge, UK), aligned to the human reference genome (hg38) using BWA (v0.7.15; Li and Durbin, 2009) and sorted and indexed using Samtools (v1.3.1). Duplicate reads are marked and removed using Picard (v2.2.4). Variant calling was performed using VarScan (v.2.4.3) with the following parameters: ≥1000<u>x</u> coverage; > <u>>1</u>000<u>x;</u> support reads, > 10≥base quality, > 30; <u>>ma</u>pping quality, > 10; >va<u>ri</u>ant threshold, < 0.05. Variant annotation was performed using VEP (v109).

### Statistical Analysis and Graphics

Data was analysed using Prism v5 using data appropriate statistical tests (detailed in the text). Statistical significance was set at p<0.05. Figures were produced using Prism v5 and BioRender (BioRender.com).

## Results

### Nanobiopsy

MtDNA from four (20%) single-cell and eleven (22%) nanobiopsy lysate samples, isolated from human skeletal muscle tissue (>1ng input; n = 1; **Figure 2**), were successfully enriched using targeted PCR and then sequenced. Nanobiopsy lysate underwent NGS (MiSeq, Illumina) with an average sequencing depth of >10,000 (**Figure 3**). This is in line with previous cell-culture based nanobiopsy experiments (39) and other subcellular isolation techniques (41,42) and compares favourably with traditional single-cell approaches such as LCM (44,52,53,54). Single-cell lysate samples were sanger sequenced alongside two positive gDNA controls.

**Figure 2.**
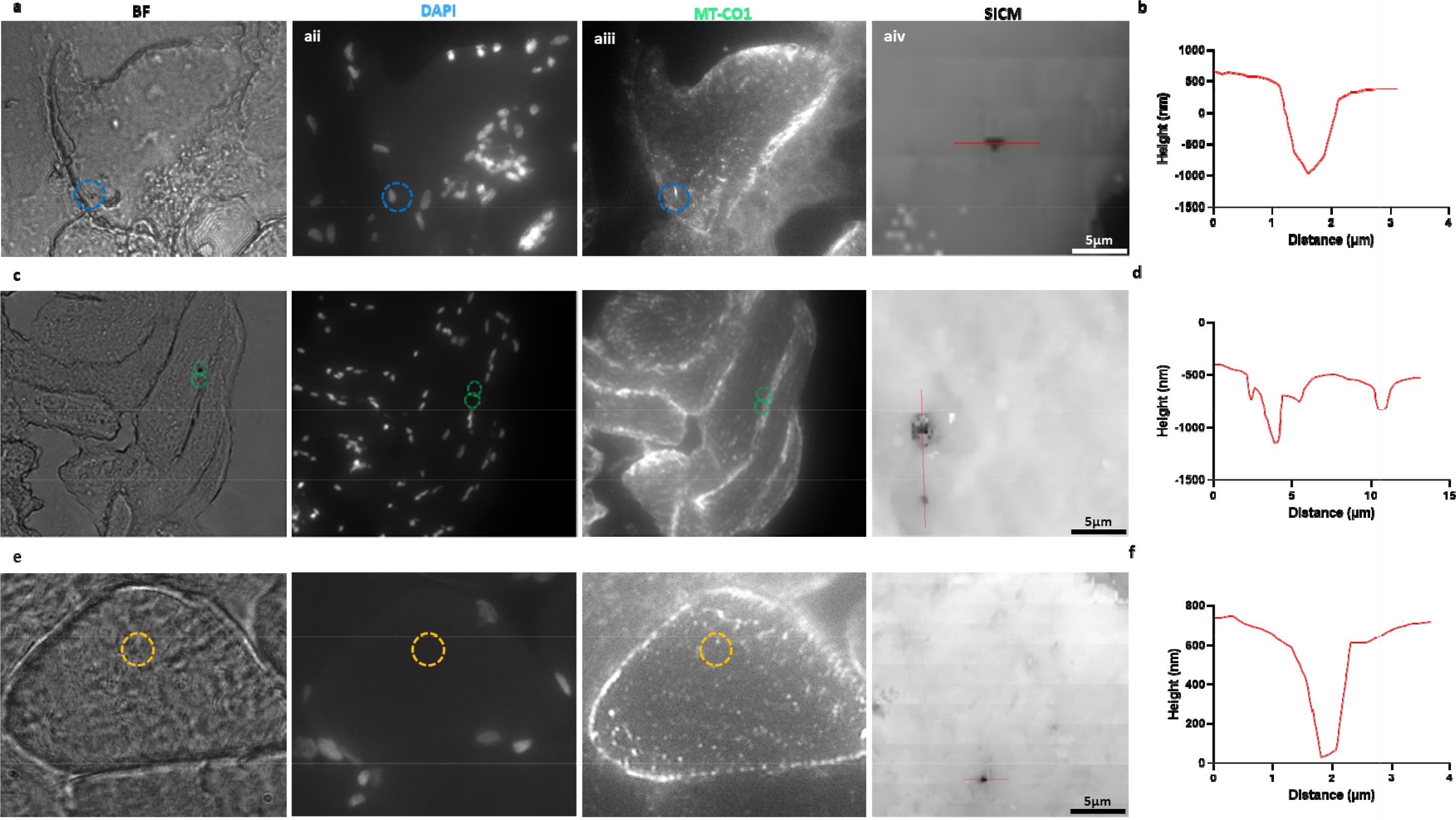
Post-biopsy images were taken from one of three distinct subcellular foci from human skeletal muscle fibres: perinuclear (PN, blue, top); subsarcolemmal (SS, green, middle) or intermyofibrillar (IMF, yellow, bottom). The PN nanobiopsy corresponds to biopsy lysate samples 22, the SS to biopsy lysate sample 25 and the IMF to biopsy lysate sample 51. Post-biopsy images were taken corresponding to bright-field (BF); DAPI nuclear stain; mitochondrial respiratory chain Complex IV, subunit I (MT-CO1) as well as SICM topographical images (a,c,e) and scans (b,d,f) were also taken.

**Figure 3.**
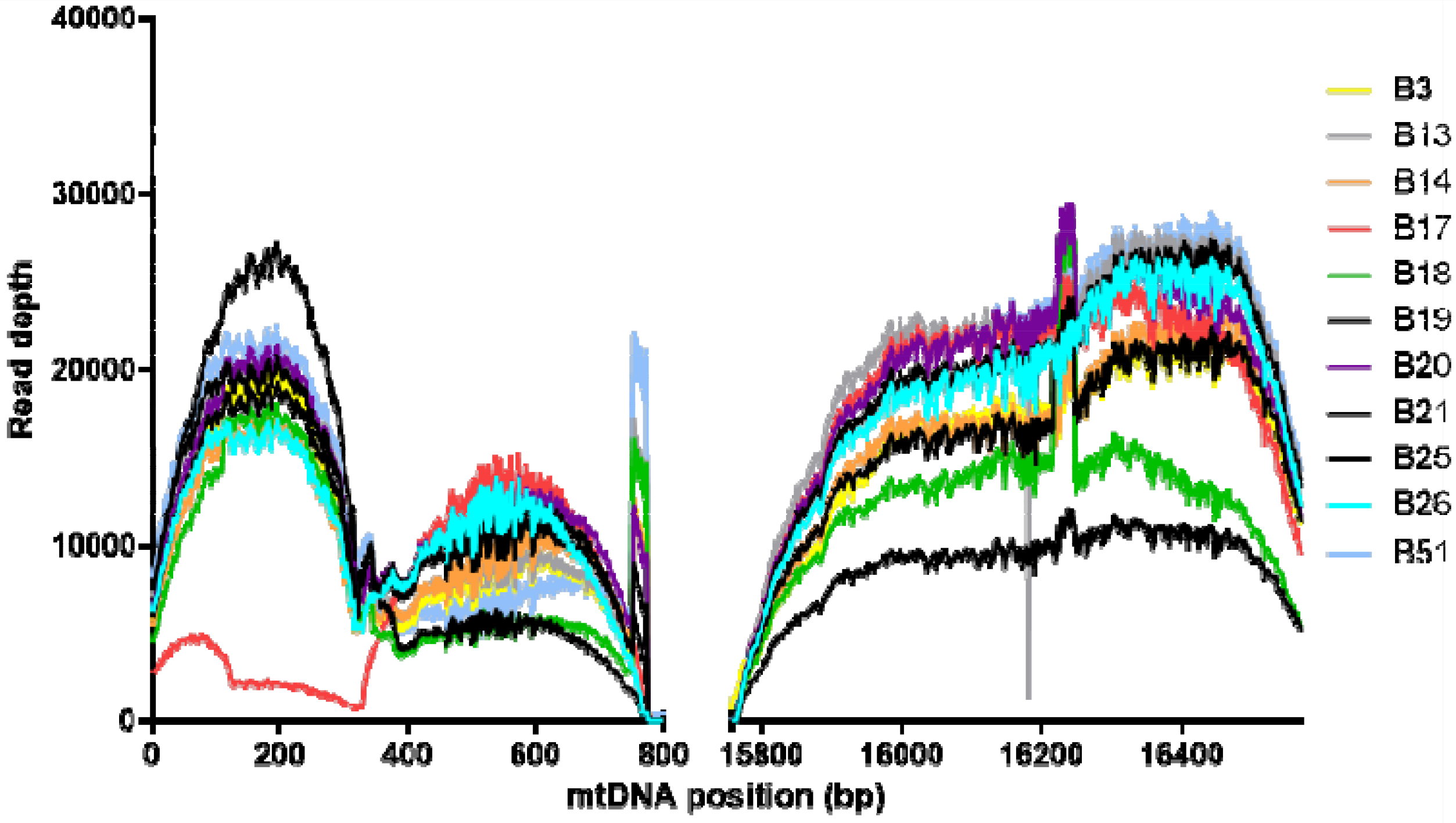
Depicted is a graph showing the breadth of coverage of Illumina MiSeq spanning the entirety of the mtDNA control region. Each plotted line corresponds to the read depth of mtDNA sequenced from individual nanobiopsy lysate samples acquired from human skeletal muscle tissue. The depth of sequencing coverage ranges from 0 to 29430. The mean depth of coverage was 10911-fold. Peaks in the read depth correspond to D2 and D4 primer binding sites at mt.750 and mt.16250 (**Table 2**).

### Subcellular mtDNA sequencing

Following NGS, 12 homoplasmic variants were detected in all nanobiopsy lysate samples compared with three homoplasmic variants in control gDNA: mt.263A>G was observed in all nanobiopsy and gDNA samples, mt.750A>G was observed in all gDNA and ten nanobiopsy lysate samples, mt.73A>G was observed in all gDNA and nine nanobiopsy lysate samples and mt.16129T>C was observed in six nanobiopsy lysate samples but neither gDNA controls (**Figure 4**). The mtSNV mt.73A>G was also shown at heteroplasmic levels in fibres 3 (86%) and 4 (81%), whilst mt.16126T>C was observed at heteroplasmic levels in fibre three: biopsy 18 (13%), fibre four: biopsies 20 (97%) and 21 (25%) and absent from fibre two . The 750A>G mtSNV was not observed in fibre 6 and mt.16126T>C was observed at only 10% (**Figure 4**). This supports the clonal expansion or loss of mutations at the subcellular level, in disparate fashion to the tissue consensus.

**Figure 4.**
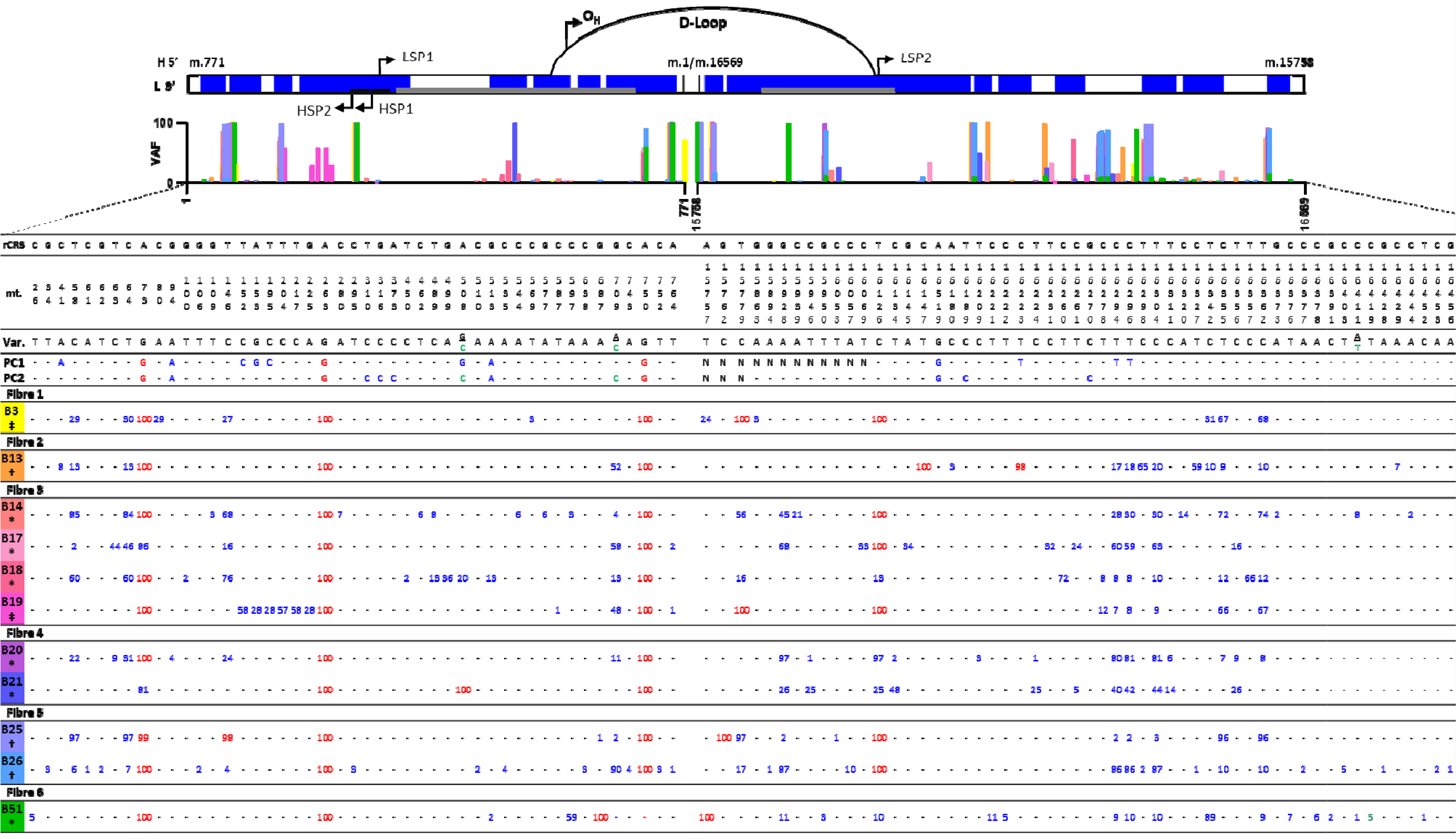
The top graphic represents all sequenced nanobiopsy variants (blue) mapped to the D-loop region of the mitochondrial genome (black). Grey bars correspond to hypervariable sections (HVS) of the mitochondrial genome: HVS1 (left) and HVS2 (right). Also depicted are the heavy strand replication origin (O_H_), as well as heavy (HSP)and light stand promoters (LSP) (56). The bar graph shows the mitochondrial variant allele frequencies (VAF, %) corresponding to colour-coded nanobiopsy lysate samples. Positive control (PC) gDNA samples were sequenced using Sanger sequencing and are represented by the substituted bases – heteroplasmic (blue) or homoplasmic (red). Nanobiopsies (B) underwent MiSeq sequencing (Illumina) and are listed by fibre and nanobiopsy lysate number - heteroplasmic (blue) or homoplasmic (red). The positions of all mitochondrial single nucleotide variants (mtSNVs) are noted relative to the rCRS (NC_012920.1). For nanobiopsy lysate samples, numbers correspond to variant allele frequency. Subcellular foci of nanobiopsies are noted as: ‡ = PN; † = SS; * = IMF. - = wild-type. N = not successfully sequenced.

A total of 103 mtDNA single nucleotide variants (mtSNVs) were detected in the D-loop of nanobiopsy lysate samples. This was a significantly greater number than gDNA variants (p< 0.01, Student’s t-test; **Figure 4**). An average of 20 mtSNVs per biopsy were sequenced, which is comparable with previous studies (55). This is compared with 21 mtSNVs in positive control gDNA. Of the 103 distinct mtSNVs from nanobiopsy lysate samples, 86 (39%) were low-level heteroplasmy (<10%) (11). Some heteroplasmies were consistent across the majority of cells e.g., mt.16519 in biopsy 13 (74%), fibre two; biopsies 17 (77%) and 18 (92%), fibre three; biopsies 20 (86%) and 21 (80%), fibre four and biopsy 26 (90%), fibre five. Some heteroplasmic variants were specific to individual cells e.g., mt.80C>A (29%) in fibre 1. Other specific variants, e.g., mt.709 and mt.16294 in fibre 3, showed evidence of expansion between and within cells, with heteroplasmy levels ranging between 4-58% and 7-60% respectively (**Figure 4**). Notably, we observed no nDNA variant detection in any biopsy.

There was a significant difference in total heteroplasmy level between all nanobiopsy lysate samples (p⍰1<⍰10.01, Kruskal-Wallis test; **Figure 5a**) but only moderate significance in heteroplasmy between individual fibres (p = 0.05, Kruskal-Wallis test; **Figure 5b**) and no significant difference between skeletal muscle fibre foci (p >0.05, Kruskal-Wallis test; **Figure 5c**). Some biopsies exhibited a wide range of heteroplasmy (biopsies 3-21; **Figure 5a**). others showed a more bi-modal distribution of heteroplasmy (biopsies 25,26,51; **Figure 5a**). This was also reflected in skeletal muscle fibres, with fibres one to four showing a greater spread of heteroplasmy and a bi-modal heteroplasmy distribution being observed in fibres five and six (**Figure 5b**).

**Figure 5.**
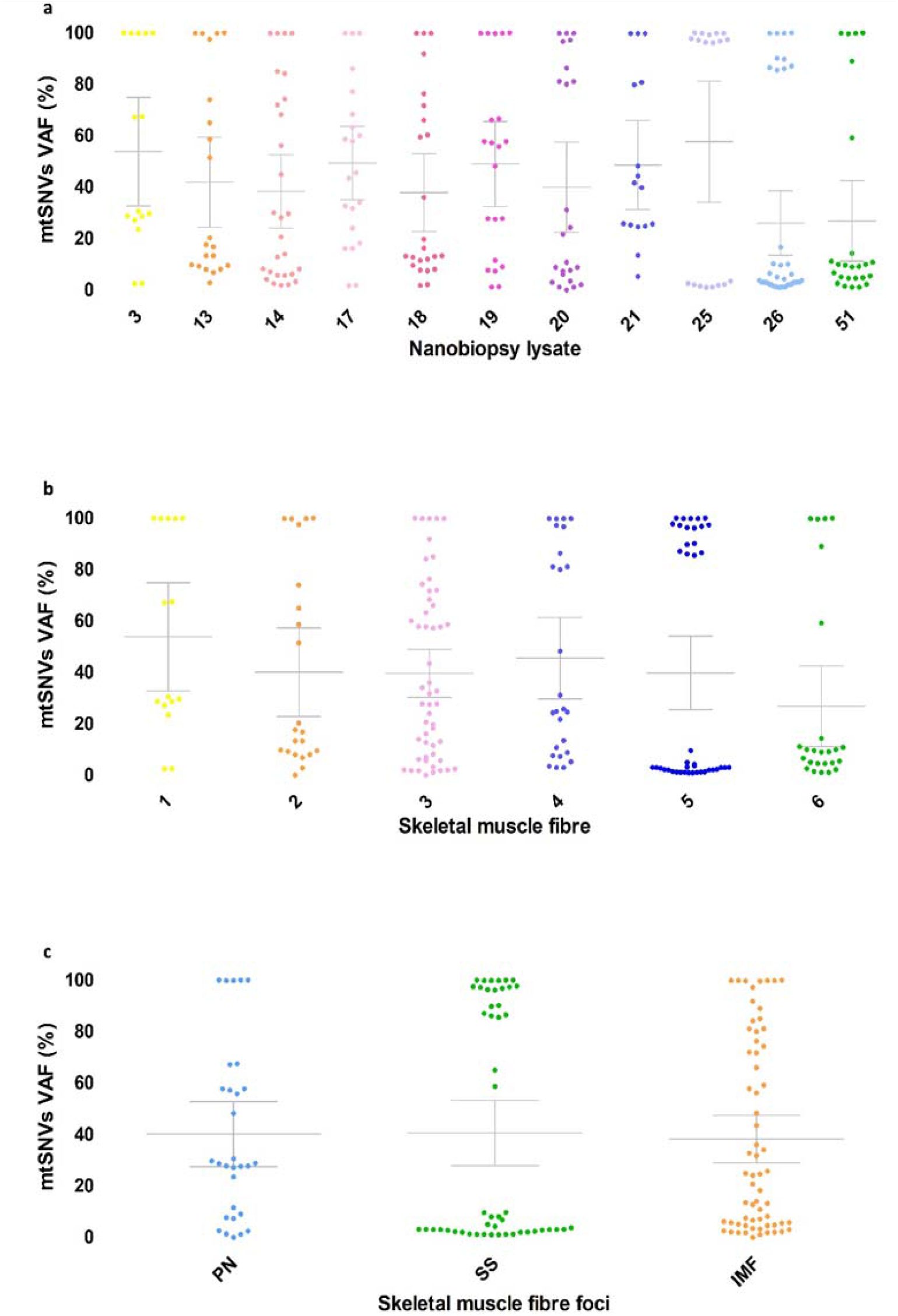
Data points represent the percentage Variable Allele Frequency (VAF) of the individual mitochondrial single nucleotide variants (mtSNVs). The error bars depict the mean allele frequency and 95% CI. (a) There was a significant difference in median heteroplasmy between individual nanobiopsy lysate samples (*p* = 0.004, Kruskal-Wallis test). (b) Although a difference in heteroplasmy between nanobiopsy samples was observed, this did not reach statistical significance (*p* = 0.054, Kruskal-Wallis test). (c) Biopsies were taken from three subcellular foci: Perinuclear (PN; *n* = 2, mean = 40.2, 95% CI⍰= ⍰27.6 – 52.9; mean median = 28.8), subsarcolemmal (SS; *n* = 3, mean = 40.6, 95% CI⍰= ⍰27.9 – 53.3; mean median = 7.5) and intermyofibrillar (IMF; *n* = 6, mean = 38.3, 95% CI⍰= ⍰29.1 – 47.5; mean median = 24.4). No significant difference in heteroplasmy between the different foci was observed (*p* = 0.555, Kruskal-Wallis test).

## Discussion

In this study we demonstrate, for the first time, the subcellular sequencing of mtDNA acquired from subcellular skeletal muscle foci using nanobiopsy in tandem with MiSeq.

A long-standing challenge in analysing low-level heteroplasmy was the ability of sequencing technologies to accurately distinguish low-level signals from noise (57,58,59). Although the advent of NGS technologies has made this possible (11), remaining challenges include the inherent noise associated with bulk population analysis when trying to disseminate single-or subcellular level heteroplasmy. Subcellular sequencing within neurons has been demonstrated, capable of detecting down to 10% heteroplasmy (55). Here we take this to the next level, in terms of sampling precision and sequencing resolution. There was a significantly greater number of successfully sequenced heteroplasmic nanobiopsy variants compared with gDNA variants. Whilst this could partly be attributed to the greater sequencing depth and efficiency of the MiSeq platform (60), the observed elevation in nanobiopsy heteroplasmy is indicative of subtle subcellular ‘microheteroplasmy’ that was previously indistinguishable from the background noise associated with the sequencing of single-cells or homogenate (24,61). Whilst very low-level nanobiopsy heteroplasmy (<1%) was detected with MiSeq, 1% heteroplasmy was set as a threshold for analysis.

Homoplasmic variants (≥98%) observed in gDNA were also observed in nanobiopsy samples and correspond to haplogroup markers (58,62), though not all sequenced homoplasmic mtSNVs were observed in gDNA samples. This was likely to have been affected by the partial sequencing of positive control D1 amplicons, which coincides with the hypervariable segment 1 (HSV1, **Figure 4**)(63).

The observed difference in heteroplasmy between biopsies is reflective of the genetic heterogeneity that exists not just between cells but within subcellular mitochondrial populations. Whilst the difference in overall heteroplasmy load, between different fibres, did not reach significance (**Figure 5b**), the specific polymorphisms and range of heteroplasmy that contribute to total heteroplasmy are different between different fibres (**Figure 4**). No significant difference was observed between heteroplasmy level between different subcellular foci; however, this could be affected by lack of biopsies where all foci were samples within the same fibre -for direct comparison (**Figure 4 and 5c**). The observed differences in heteroplasmy distribution between different biopsies, foci, and fibres, whilst exciting, cannot yet be fully explained. Different patterns of heteroplasmy could reflect different mutation thresholds (12,20). This may also be reflective of stochastic or selective clonal expansion events (18).

Having validated the nanobiopsy technique to investigate subcellular heteroplasmy from distinct foci, an obvious next step is to investigate localised patterns in subcellular heteroplasmy within diseased tissues to elucidate specific mechanisms responsible for clonal expansion of mtDNA mutations (18,29). Our approach allows the investigation of subcellular heteroplasmy to other tissues and cultured cells as means of investigating spatial and temporal patterns in mtDNA mutation distribution. Recent advancements in Oxford Nanopore Sequencing technologies also offer the future potential for enrichment free ‘pore-to-pore’ sequencing (64). Nanobiopsy in combination with existing and emerging sequencing technologies will allow us to harness the untapped potential of these platforms to perform subcellular sequencing (61), this is increasingly vital as we move away from the era of single-cell omics towards subcellular-omics (65).

## Acknowledgements

This work was supported by a Wellcome Trust grant to the Wellcome Centre for Mitochondrial Research [203105]. A.B. acknowledges funding from the MRC DiMeN Doctoral Training Partnership [OSR/0200/2018/BURY]. A.E.V. acknowledges funding through a Sir Henry Wellcome Postdoctoral Fellowship [215888, https://doi.org/10.35802/215888]. P.A. acknowledges funding from the EPSRC [EP/S01764x/1].

## References

1. Schon EA, DiMauro S, Hirano M. Human mitochondrial DNA: roles of inherited and somatic mutations. Nature Reviews Genetics. 2012;13(12):878–90.

2. Skladal D, Sudmeier C, Konstantopoulou V, Stöckler-Ipsiroglu S, Plecko-Startinig B, Bernert G, et al. The clinical spectrum of mitochondrial disease in 75 pediatric patients. Clinical pediatrics. 2003;42(8):703–10.

3. Gorman GS, Schaefer AM, Ng Y, Gomez N, Blakely EL, Alston CL, et al. Prevalence of nuclear and mitochondrial DNA mutations related to adult mitochondrial disease. Annals of neurology. 2015;77(5):753–9.

4. Alston CL, Rocha MC, Lax NZ, Turnbull DM, Taylor RW. The genetics and pathology of mitochondrial disease. The Journal of pathology. 2017;241(2):236–50.

5. Bender A, Krishnan KJ, Morris CM, Taylor GA, Reeve AK, Perry RH, et al. High levels of mitochondrial DNA deletions in substantia nigra neurons in aging and Parkinson disease. Nature genetics. 2006;38(5):515–7.

6. Coxhead J, Kurzawa-Akanbi M, Hussain R, Pyle A, Chinnery P, Hudson G. Somatic mtDNA variation is an important component of Parkinson’s disease. Neurobiology of aging. 2016;38:217. e1-. e6.

7. Bury AG, Pyle A, Elson JL, Greaves L, Morris CM, Hudson G, et al. Mitochondrial DNA changes in pedunculopontine cholinergic neurons in Parkinson disease. Annals of neurology. 2017;82(6):1016–21.

8. Krishnan KJ, Ratnaike TE, De Gruyter HL, Jaros E, Turnbull DM. Mitochondrial DNA deletions cause the biochemical defect observed in Alzheimer’s disease. Neurobiology of aging. 2012;33(9):2210–4.

9. Yu-Wai-Man P, Lai-Cheong J, Borthwick GM, He L, Taylor GA, Greaves LC, et al. Somatic mitochondrial DNA deletions accumulate to high levels in aging human extraocular muscles. Investigative ophthalmology & visual science. 2010;51(7):3347–53.

10. Greaves LC, Nooteboom M, Elson JL, Tuppen HA, Taylor GA, Commane DM, et al. Clonal expansion of early to mid-life mitochondrial DNA point mutations drives mitochondrial dysfunction during human ageing. PLoS genetics. 2014;10(9):e1004620.

11. Payne BA, Wilson IJ, Yu-Wai-Man P, Coxhead J, Deehan D, Horvath R, et al. Universal heteroplasmy of human mitochondrial DNA. Human molecular genetics. 2013;22(2):384–90.

12. Rossignol R, Faustin B, Rocher C, Malgat M, Mazat J-P, Letellier T. Mitochondrial threshold effects. Biochemical Journal. 2003;370(3):751–62.

13. Taylor RW, Barron MJ, Borthwick GM, Gospel A, Chinnery PF, Samuels DC, et al. Mitochondrial DNA mutations in human colonic crypt stem cells. The Journal of clinical investigation. 2003;112(9):1351–60.

14. Trifunovic A, Wredenberg A, Falkenberg M, Spelbrink JN, Rovio AT, Bruder CE, et al. Premature ageing in mice expressing defective mitochondrial DNA polymerase. Nature. 2004;429(6990):417–23.

15. Ross JM, Stewart JB, Hagström E, Brené S, Mourier A, Coppotelli G, et al. Germline mitochondrial DNA mutations aggravate ageing and can impair brain development. Nature. 2013;501(7467):412–5.

16. Nekhaeva E, Bodyak ND, Kraytsberg Y, McGrath SB, Van Orsouw NJ, Pluzhnikov A, et al. Clonally expanded mtDNA point mutations are abundant in individual cells of human tissues. Proceedings of the National Academy of Sciences. 2002;99(8):5521–6.

17. Pinto M, Moraes CT. Mechanisms linking mtDNA damage and aging. Free Radical Biology and Medicine. 2015;85:250–8.

18. Lawless C, Greaves L, Reeve AK, Turnbull DM, Vincent AE. The rise and rise of mitochondrial DNA mutations. Open biology. 2020;10(5):200061.

19. Baines HL, Stewart JB, Stamp C, Zupanic A, Kirkwood TB, Larsson N-G, et al. Similar patterns of clonally expanded somatic mtDNA mutations in the colon of heterozygous mtDNA mutator mice and ageing humans. Mechanisms of ageing and development. 2014;139:22–30.

20. Stewart JB, Chinnery PF. Extreme heterogeneity of human mitochondrial DNA from organelles to populations. Nature Reviews Genetics. 2021;22(2):106–18.

21. Ahmed ST, Craven L, Russell OM, Turnbull DM, Vincent AE. Diagnosis and treatment of mitochondrial myopathies. Neurotherapeutics. 2018;15:943–53.

22. Grady JP, Pickett SJ, Ng YS, Alston CL, Blakely EL, Hardy SA, et al. mt DNA heteroplasmy level and copy number indicate disease burden in m. 3243A>G mitochondrial disease. EMBO molecular medicine. 2018;10(6):e8262.

23. Anand RK, Chiu DT. Analytical tools for characterizing heterogeneity in organelle content. Current opinion in chemical biology. 2012;16(3-4):391-9.

24. Aryaman J, Johnston IG, Jones NS. Mitochondrial heterogeneity. Frontiers in genetics. 2019;9:718.

25. Ngo J, Osto C, Villalobos F, Shirihai OS. Mitochondrial heterogeneity in metabolic diseases. Biology. 2021;10(9):927.

26. Pereira CV, Gitschlag BL, Patel MR. Cellular mechanisms of mtDNA heteroplasmy dynamics. Critical reviews in biochemistry and molecular biology. 2021;56(5):510–25.

27. Jackson CB, Turnbull DM, Minczuk M, Gammage PA. Therapeutic manipulation of mtDNA heteroplasmy: a shifting perspective. Trends in molecular medicine. 2020;26(7):698–709.

28. van den Ameele J, Li AY, Ma H, Chinnery PF, editors. Mitochondrial heteroplasmy beyond the oocyte bottleneck. Seminars in Cell & Developmental Biology; 2020: Elsevier.

29. Vincent AE, Rosa HS, Pabis K, Lawless C, Chen C, Grünewald A, et al. Subcellular origin of mitochondrial DNA deletions in human skeletal muscle. Annals of neurology. 2018;84(2):289–301.

30. Insalata F, Hoitzing H, Aryaman J, Jones NS. Stochastic survival of the densest and mitochondrial DNA clonal expansion in aging. Proceedings of the National Academy of Sciences. 2022;119(49):e2122073119.

31. Coller HA, Khrapko K, Bodyak ND, Nekhaeva E, Herrero-Jimenez P, Thilly WG. High frequency of homoplasmic mitochondrial DNA mutations in human tumors can be explained without selection. Nature genetics. 2001;28(2):147–50.

32. Elson J, Samuels D, Turnbull D, Chinnery P. Random intracellular drift explains the clonal expansion of mitochondrial DNA mutations with age. The American Journal of Human Genetics. 2001;68(3):802–6.

33. Johnston IG, Burgstaller JP, Havlicek V, Kolbe T, Rülicke T, Brem G, et al. Stochastic modelling, Bayesian inference, and new in vivo measurements elucidate the debated mtDNA bottleneck mechanism. Elife. 2015;4:e07464.

34. Burgstaller JP, Johnston IG, Poulton J. Mitochondrial DNA disease and developmental implications for reproductive strategies. Molecular human reproduction. 2015;21(1):11–22.

35. Vincent AE, White K, Davey T, Philips J, Ogden RT, Lawless C, et al. Quantitative 3D mapping of the human skeletal muscle mitochondrial network. Cell reports. 2019;26(4):996-1009. e4.

36. Campbell G, Krishnan KJ, Deschauer M, Taylor RW, Turnbull DM. Dissecting the mechanisms underlying the accumulation of mitochondrial DNA deletions in human skeletal muscle. Human molecular genetics. 2014;23(17):4612–20.

37. Shakoor A, Gao W, Zhao L, Jiang Z, Sun D. Advanced tools and methods for single-cell surgery. Microsystems & Nanoengineering. 2022;8(1):47.

38. Unwin P. Concluding remarks: next generation nanoelectrochemistry–next generation nanoelectrochemists. Faraday Discussions. 2022;233:374–91.

39. Actis P, Maalouf MM, Kim HJ, Lohith A, Vilozny B, Seger RA, et al. Compartmental genomics in living cells revealed by single-cell nanobiopsy. ACS nano. 2014;8(1):546–53.

40. Actis P. Sampling from single cells. Small Methods. 2018;2(3):1700300.

41. Nadappuram BP, Cadinu P, Barik A, Ainscough AJ, Devine MJ, Kang M, et al. Nanoscale tweezers for single-cell biopsies. Nature nanotechnology. 2019;14(1):80–8.

42. Chen W, Guillaume-Gentil O, Rainer PY, Gäbelein CG, Saelens W, Gardeux V, et al. Live-seq enables temporal transcriptomic recording of single cells. Nature. 2022;608(7924):733–40.

43. Elnathan R, Barbato MG, Guo X, Mariano A, Wang Z, Santoro F, et al. Biointerface design for vertical nanoprobes. Nature Reviews Materials. 2022:1–21.

44. Bury AG, Pyle A, Marcuccio F, Turnbull DM, Vincent AE, Hudson G, et al. A subcellular cookie cutter for spatial genomics in human tissue. Analytical and Bioanalytical Chemistry. 2022;414(18):5483–92.

45. Novak P, Li C, Shevchuk AI, Stepanyan R, Caldwell M, Hughes S, et al. Nanoscale live-cell imaging using hopping probe ion conductance microscopy. Nature methods. 2009;6(4):279–81.

46. Xu X, Valavanis D, Ciocci P, Confederat S, Marcuccio F, Lemineur J-F, et al. The new era of high-throughput nanoelectrochemistry. Analytical Chemistry. 2023;95(1):319–56.

47. Wei W, Schon KR, Elgar G, Orioli A, Tanguy M, Giess A, et al. Nuclear-embedded mitochondrial DNA sequences in 66,083 human genomes. Nature. 2022;611(7934):105–14.

48. Rocha MC, Grady JP, Grünewald A, Vincent A, Dobson PF, Taylor RW, et al. A novel immunofluorescent assay to investigate oxidative phosphorylation deficiency in mitochondrial myopathy: understanding mechanisms and improving diagnosis. Scientific reports. 2015;5(1):15037.

49. Lowes H, Pyle A, Duddy M, Hudson G. Cell-free mitochondrial DNA in progressive multiple sclerosis. Mitochondrion. 2019;46:307–12.

50. Trifunov S, Pyle A, Valentino ML, Liguori R, Yu-Wai-Man P, Burté F, et al. Clonal expansion of mtDNA deletions: different disease models assessed by digital droplet PCR in single muscle cells. Scientific reports. 2018;8(1):11682.

51. Bury AG, Robertson FM, Pyle A, Payne BA, Hudson G. The isolation and deep sequencing of mitochondrial DNA. Mitochondrial Medicine: Volume 3: Manipulating Mitochondria and Disease-Specific Approaches: Springer; 2021. p. 433–47.

52. Cantuti-Castelvetri I, Lin MT, Zheng K, Keller-McGandy CE, Betensky RA, Johns DR, et al. Somatic mitochondrial DNA mutations in single neurons and glia. Neurobiology of aging. 2005;26(10):1343–55.

53. Pflugradt R, Schmidt U, Landenberger B, Sänger T, Lutz-Bonengel S. A novel and effective separation method for single mitochondria analysis. Mitochondrion. 2011;11(2):308–14.

54. Lin MT, Cantuti-Castelvetri I, Zheng K, Jackson KE, Tan YB, Arzberger T, et al. Somatic mitochondrial DNA mutations in early Parkinson and incidental Lewy body disease. Annals of neurology. 2012;71(6):850–4.

55. Morris J, Na Y-J, Zhu H, Lee J-H, Giang H, Ulyanova AV, et al. Pervasive within-mitochondrion single-nucleotide variant heteroplasmy as revealed by single-mitochondrion sequencing. Cell reports. 2017;21(10):2706–13.

56. Tan BG, Mutti CD, Shi Y, Xie X, Zhu X, Silva-Pinheiro P, et al. The human mitochondrial genome contains a second light strand promoter. Molecular Cell. 2022;82(19):3646-60. e9.

57. White HE, Durston VJ, Seller A, Fratter C, Harvey JF, Cross NC. Accurate detection and quantitation of heteroplasmic mitochondrial point mutations by pyrosequencing. Genetic testing. 2005;9(3):190–9.

58. Elliott HR, Samuels DC, Eden JA, Relton CL, Chinnery PF. Pathogenic mitochondrial DNA mutations are common in the general population. The American journal of human genetics. 2008;83(2):254–60.

59. Fazzini F, Fendt L, Schönherr S, Forer L, Schöpf B, Streiter G, et al. Analyzing low-level MtDNA heteroplasmy—Pitfalls and challenges from bench to benchmarking. International Journal of Molecular Sciences. 2021;22(2):935.

60. Guo Y, Li C-I, Sheng Q, Winther JF, Cai Q, Boice JD, et al. Very low-level heteroplasmy mtDNA variations are inherited in humans. Journal of genetics and genomics. 2013;40(12):607–15.

61. Duan M, Tu J, Lu Z. Recent advances in detecting mitochondrial DNA heteroplasmic variations. Molecules. 2018;23(2):323.

62. Yonova-Doing E, Calabrese C, Gomez-Duran A, Schon K, Wei W, Karthikeyan S, et al. An atlas of mitochondrial DNA genotype–phenotype associations in the UK Biobank. Nature genetics. 2021;53(7):982–93.

63. Wei W, Tuna S, Keogh MJ, Smith KR, Aitman TJ, Beales PL, et al. Germline selection shapes human mitochondrial DNA diversity. Science. 2019;364(6442):eaau6520.

64. Zascavage RR, Thorson K, Planz JV. Nanopore sequencing: An enrichment-free alternative to mitochondrial DNA sequencing. Electrophoresis. 2019;40(2):272–80.

65. Tharkeshwar AK, Gevaert K, Annaert W. Organellar omics—a reviving strategy to untangle the biomolecular complexity of the cell. Proteomics. 2018;18(5-6):1700113.

